# Cholesterol bound *Plasmodium falciparum* co-chaperone ‘PFA0660w’ complexes with major virulence factor ‘PfEMP1’ via chaperone ‘PfHsp70-x’

**DOI:** 10.1101/311134

**Authors:** Ankita Behl, Vikash Kumar, Anjali Bisht, Jiban J. Panda, Rachna Hora, Prakash Chandra Mishra

**Author notes:** Corresponding author name: Prakash Chandra Mishra; Extn: 3224.

## Abstract

Lethality of *Plasmodium falciparum* (Pf) caused malaria results from ‘cytoadherence’, which is effected by exported *Plasmodium falciparum* erythrocyte membrane protein 1 (PfEMP1) family. Several exported Pf proteins (exportome) including chaperones alongside cholesterol rich microdomains are crucial for PfEMP1 translocation to infected erythrocyte surface. An exported Hsp40 (heat shock protein 40) ‘PFA0660w’ functions as a co-chaperone of ‘PfHsp70-x’, and these co-localize to specialized intracellular mobile structures termed J-dots. Our studies attempt to understand the function of PFA0660w-PfHsp70-x chaperone pair using recombinant proteins. Biochemical assays reveal that N and C-terminal domains of PFA0660w and PfHsp70-x respectively are critical for their activity. We show the novel direct interaction of PfHsp70-x with the cytoplasmic tail of PfEMP1, and binding of PFA0660w with cholesterol. PFA0660w operates both as a chaperone and lipid binding molecule via its separate substrate and cholesterol binding sites. PfHsp70-x binds cholesterol linked PFA0660w and PfEMP1 simultaneously *in vitro* to form a complex. Collectively, our results and the past literature support the hypothesis that PFA0660w-PfHsp70-x chaperone pair assists PfEMP1 transport across the host erythrocyte through cholesterol containing ‘J-dots’. Since PFA0660w seems essential for parasite survival, characterization of its interaction with PfHsp70-x and J-dots may form the basis for development of future antimalarials.

## Introduction

*P. falciparum* is the major cause of severe complicated malaria that is responsible for ~0.4 million deaths annually ^1^. This high mortality rate results from the ability of iRBCs to bind host endothelial receptors by the process of cytoadherence, which is largely due to the PfEMP1 family of proteins ^2,3^ PfEMP1 is one of the several exported protein families of Pf that play a crucial role in host erythrocyte remodelling to facilitate virulence, growth and survival of the parasites ^4^. Most exported proteins carry a PEXEL (*Plasmodium* export element) motif ^5^, while a few are exported in the absence of this signal ^6^. Post infection, the parasite establishes its own protein trafficking machinery including sub-cellular organelles like Maurer’s clefts (MCs) ^7^ etc. to facilitate protein export. PfEMP1 export has been linked to several proteins like PfSBP1, MAHRP, PfEMP3 ^8–10^ and various chaperones ^11^ that support transport of this major virulence antigen across the parasite confines.

Inside the cellular environment, chaperones prevent aggregation or misfolding of nascent polypeptides, hence accomplishing the conformational integrity of the entire proteome. The Pf genome encodes numerous chaperones including members of heat shock proteins of ~40kDa, ~ 60kDa, ~70kDa and ~90kDa families that are upregulated in response to stress caused by parasite infection ^12–16^. Amongst all chaperones, Hsp40 members are an abundant class distributed into four types (I to IV) ^17^. These members of the ‘malaria exportome’ ^5^ play crucial roles in cellular processes like protein translation, folding, translocation, and degradation ^17^, underscoring their importance in parasite biology. Hsp40s are molecular co-chaperones of Hsp70s, and contain a highly conserved J-domain ^18^. They target protein substrates to Hsp70s for folding, and stabilize them in the substrate bound form ^19–21^. The J domain exhibits four helices and a highly conserved His-Pro-Asp (HPD) tripeptide motif that is vital for stimulating the ATPase activity of Hsp70 ^22,23^. Hsp70 proteins consist of two distinct functional domains viz. a 45 kDa N-terminal ATPase domain, and a 25 kDa C-terminal substrate binding domain (SBD). The SBD acts as lid and facilitates entrapment of the substrate ^24,25^.

MCs have been long considered as intermediary compartments for PfEMP1 export ^26,27^. More recently, Kulzer *et al*., reported cholesterol containing novel mobile structures termed ‘J-dots’ to be involved in trafficking of parasite encoded proteins (e.g. PfEMP1) across the host cytosol ^28^. Furthermore, delivery of this key protein is reported to involve its insertion at cholesterol rich membrane microdomains ^29^. Chaperone complexes from other species have also been shown to associate with detergent resistant lipid rafts rich in cholesterol and sphingolipids, possibly through Hsp70 ^30,31^. J dots are known to carry exported PfHsp40 chaperones (PFA0660w, PFE0055c) and PfHsp70-x present as large complexes with several other exported proteins ^28,32,33^. PFA0660w, a type II PEXEL positive PfHsp40 has been reported to be essential for survival of parasites inside erythrocytes ^5^. Immunofluorescence assays revealed partial co-localization of these GFP-tagged PfHsp40s and PfHsp70-x with PfEMP1, suggesting a role for these chaperones in PfEMP1 transport ^30,31^. This was supported by another report where deletion of PfHsp70-x led to delayed export of PfEMP1 to the erythrocyte surface ^34^. Another study elucidated the trafficking of PfHsp70-x and demonstrated that its export takes place via the PTEX translocon and is directed by an N-terminal secretory signal sequence ^35^. Though the role of PfHsp70-x and cholesterol has been implicated in export of the PfEMP1 family of proteins through *in vivo* experiments, the underlying mechanism of this process remains ambiguous.

In the present study, we provide the first direct evidence for the molecular interplay of events in this crucial process using recombinant proteins and their lipid interactions. We have performed domain characterization of the PFA0660w-PfHsp70-x chaperone pair that sheds light on how these partners interact with each other to execute their function. We also depict binding of PfHsp70-x with ‘PfEMP1’, and display cholesterol binding properties of PFA0660w. Our results show that cholesterol and substrate binding sites on PFA0660w exist in distinct pockets on the protein, highlighting its dual functionality. Our complex binding assays clearly illustrate that PfHsp70-x interacts together with both cholesterol-linked PFA0660w and PfEMP1 to form a complex. Together, our results provide mechanistic insights into the functional role of the PFA0660w-PfHsp70-x chaperone pair. These studies hence serve to fill the key knowledge gaps in the understanding of Pf biology and are very likely to form the framework for structure-based drug design against fatal malaria.

## Results

### Cloning, expression and purification of recombinant proteins

PFA0660w displays a Sis1/Hdj1 domain organization containing a J domain (encompassing the Hsp70 interaction site at the N-terminus) followed by a G/F motif rich region and a substrate binding domain (Fig. 1A, top left panel). Residues ‘RCLAE’ (58-62) represent the PEXEL motif responsible for the export of this protein to the host cell (Fig.1B). PfHsp70-x possesses two domains viz. an N-terminal nucleotide-binding domain (NBD) with ATPase activity, which is connected by a linker region to a SBD present in the C-terminal region (Fig. 1A, top right panel). Two deletion constructs each of PFA0660w (PFA0660w-C, PfHsp70-x-S) and PfHsp70-x (PfHsp70-x-S, PfHsp70-C) were cloned in T7 promoter based plasmid pET-28a(+), and expressed in the soluble form in *E. coli* BL21 (DE3) cells with a 6X hexahistidine tag (Fig. 1A, lower panels). PFA0660w-C (81-386) contains the complete conserved region of the protein including its J domain, G/F region and C-terminally located SBD, whereas PFA0660w-S (219-386) harbors only the C-terminal region with its SBD. PfHsp70-x-S (30-412) contains the N-terminal region of the protein, with its ATPase domain, whereas PfHsp70-x-C (27-679) includes both the ATPase domain and the C-terminal region ending with the ‘EEVN’ motif.

**Fig. 1.**
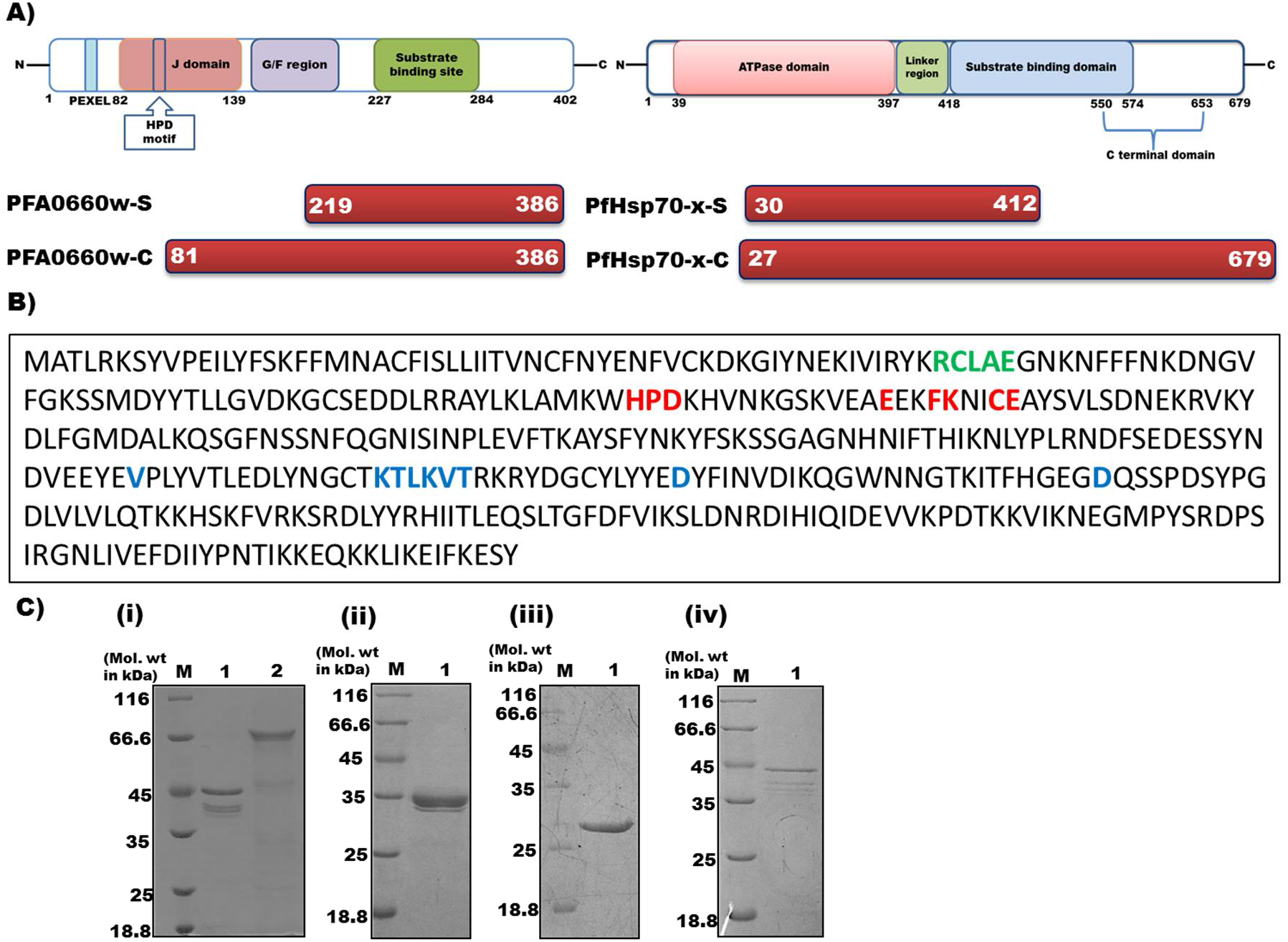
*Domain organization and expression of recombinant constructs of PFA0660w and PfHsp70-x*. (**A**) *Schematic representation of predicted domains, motifs (top panel) and cloned constructs (bottom panel) of PFA0660w (left panel) and PfHsp70-x (right panel)*. (**B**) *Protein sequence of PFA0660w with predicted features:* Residues in red denote the amino acids involved in Hsp70 interaction, blue are involved in substrate binding and green represent the PEXEL motif. (**C**) *SDS-PAGE showing recombinant purified proteins*. (i) Lane M: molecular weight marker, Lane 1: PfHsp70-x-S, Lane 2: PfHsp70-x-C. (ii) Lane M: molecular weight marker, Lane 1: PFA0660w-C. (iii) Lane M: molecular weight marker, Lane 1: PFA0660w-S. (iv) Lane M: molecular weight marker, Lane 1: ATS.

Recombinant Hsp40 and Hsp70 proteins were purified using Ni-NTA affinity and gel permeation chromatographies, while acidic terminal segment (ATS) domain of PfEMP1 was purified as reported earlier ^36,37^ Purified PfHsp70-x-S, PfHsp70-x-C, PFA0660w-C, PFA0660w-S and ATS run as species of approximately 45 kDa, 70 kDa, 38 kDa, 28 kDa and 45 kDa respectively on SDS-PAGE (Fig. 1C). Bands below the expected molecular weight can be observed particularly for PfHsp70-x-S, PfHsp70-x-C and ATS (Fig. 1C) owing to protein degradation. This is evident from western blots of recombinant proteins probed with anti-hexahistidine monoclonal antibodies (ATS^36^, Fig. S1A for Hsp40 and Hsp70 constructs). Polyclonal antisera were raised against PFA0660w-S commercially in rabbits, and specificity validated by testing on a crude extract of *E. coli* BL21 (DE3) transformed with its cloned plasmid (Fig. S1B).

### Recombinant PFA0660w and PfHsp70-x exist as monomers in solution

Dimeric Hsp40s alongwith monomeric Hsp70s have been reported to perform chaperone activity in the past ^38,39^. However, elution profiles of PFA0660w-C and PfHsp70-x-C on gel filtration column suggested that both these recombinant proteins exist as monomers (~ 38kDa and 70kDa respectively) in solution (Fig. 2A). Elution volumes of standards (BSA, chymotrypsin and lysozyme) were used for comparison to deduce the oligomeric state of these proteins (Fig. S2).

We have used *in silico* tools to structurally understand the reason for existence of PFA0660w in a monomeric form. Structures of human DnaJ subfamily B member 12 (PDB ID: 2CTP, 54% identity) and dimerization domain of *Cryptosporidium parvum* Hsp40 (PDB ID: 2Q2G, 49% identity) were used as templates for modelling J domain (78-154) and C-terminal region (226-398) of PFA0660w respectively using Swiss model tool, and refined using 3D refine (Fig. 2B). Various tests like Ramachandran plot, Verify 3D and Errat were run on the generated models to assess their acceptability, and were found suitable for structural analysis (Fig. S3).

**Fig. 2.**
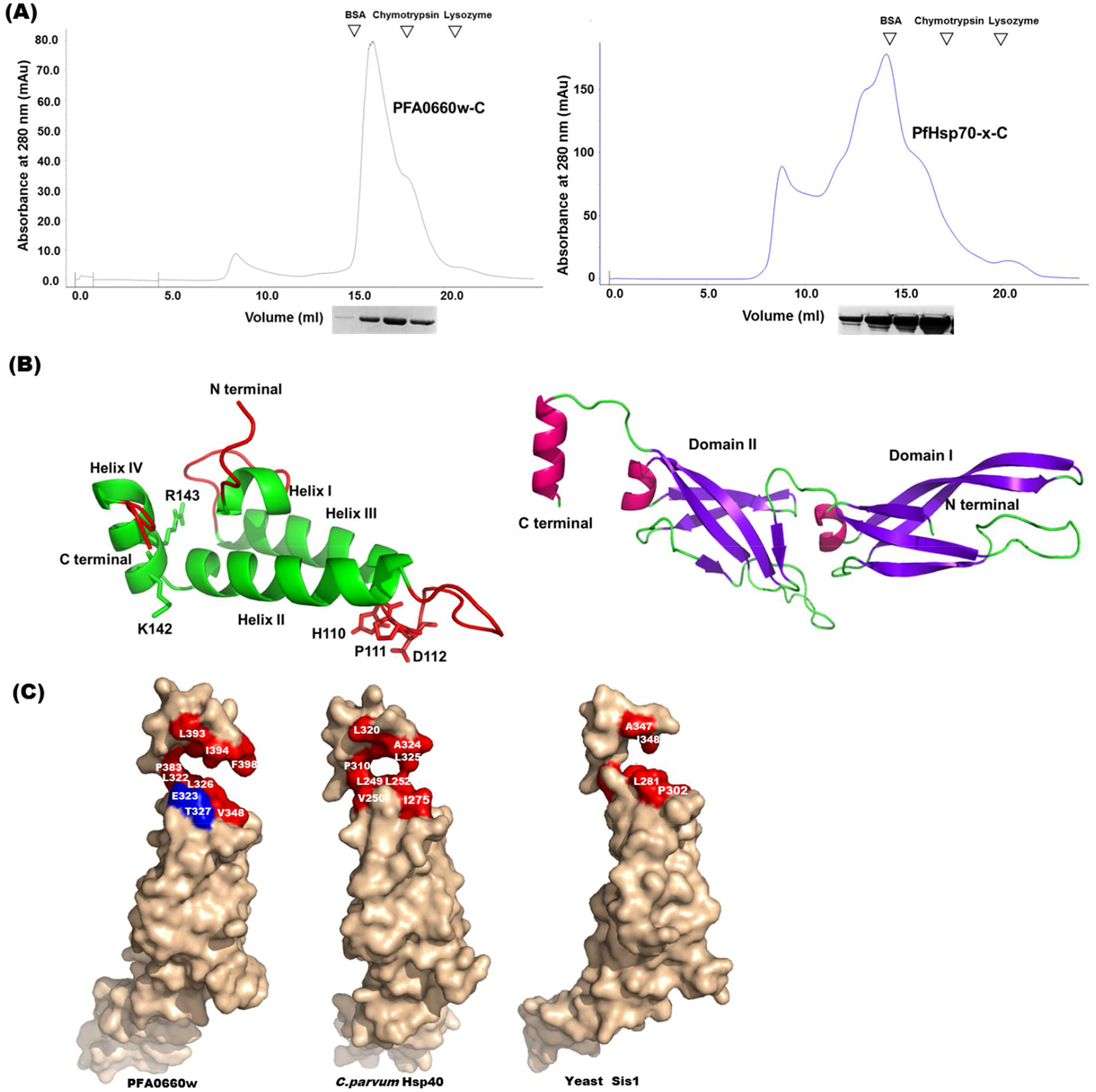
*Oligomeric state of PFA0660w and PfHsp70-x*. (**A**) *Elution profiles of PFA0660w-C and PfHsp70-x-C on gel permeation column superdex 200 10/300 GL*. SDS-PAGE of elutes is shown below the chromatogram at corresponding retention volume. Elution volume of standards on the same column are indicated by arrows (BSA monomer; 66 kDa, Chymotrypsin 25 kDa, and Lysozyme; 14.3 kDa). (**B**) *Cartoon representation of homology models of N-terminal J domain (78-154) (left panel) constituting the Hsp70 interaction site and C-terminal region (226-398) (right panel) containing the substrate binding site*. Residues involved in Hsp70 interaction are shown as ball and sticks, and are labelled in the left panel. (**C**) *Surface representation of C terminal regions of PFA0660w, C. parvum Hsp40 and yeast Sis1*. Hyrophobic residues predicted to be present at the dimer interface are shown in red, and are labelled. E323 and T327 (blue) on PFA0660w lack hydrophobicity, and correspond to V250 of *C. parvum* and L281 of Sis1 respectively. Hydrophobic dimer interface residues of PFA0660w (P351, I380, Y382, I386, I397), *C. parvum* Hsp40 (A252, P278, I307, F309, L313, I321) and yeast Sis1 (F276, L280, P305, V334, Y336, P337) are obscured from view in the depicted orientation.

Atomic resolution structures of C-terminal peptide-binding fragments of type II Hsp40s from *S. cerevisiae* (Sis1, PDB ID: 1C3G) and *C. parvum* (PDB ID: 2Q2G) revealed that both exist as homo-dimers ^38,40^. The Protein interaction calculator (PIC) computed several hydrophobic interactions and hydrogen bonds to be involved in dimerization of these proteins (Table S1). Although most of these interactions were found conserved in our modelled structure, a few hydrophobic interactions were found absent from the dimer interface. Specifically, V250 (helix II, chain A) - L325 (helix III, chain B) and A252 (helix II, chain A) – F309 (helix III, chain B) hydrophobic bonds in *C. parvum* Hsp40 were unable to form in PFA0660w due to replacement of non-polar residues V250 and A252 with polar amino-acids E323 and S325 respectively (Fig 2C). Also, substitution of L281 (Sis1, chain A) with T327 in PFA0660w may lead to disruption of L281 – L340 (chain B) interaction (Fig. 2C).

Additionally, *Sha et al*. have reported several residues from crystal structure of yeast Hsp40 ‘Sis1’ to form its dimer interface ^38^. These include F276, K277, L280, Y336, L340, I348 and D349 where K277 (helix II, A chain) - D349 (helix III, B chain) form a salt bridge, while F276, L280, Y336, L340 and I348 stabilise the dimer formation ^38^. Multiple sequence alignment of different type II Hsp40s showed most of these residues to be semi-conserved (Fig. S4).

### N-terminal region of PFA0660w interacts with PfHsp70-x

Binding studies of different deletion constructs using dot blot assays showed interaction of PFA0660w-C with PfHsp70-x-C and PfHsp70-x-S, (Fig. 3 A, upper panel), whereas PFA0660w-S was unable to bind either PfHsp70-x construct (Fig. 3 A, lower panel). Integrated density analysis of each dot demonstrated significant binding of PFA0660w-C with constructs of PfHsp70-x (p<0.05) (Fig. 3A, upper panel). Deletion constructs of PFA0660w did not bind BSA (negative control), which depicts the specificity of the assay. Inability of PFA0660w-S to bind PfHsp70-x also behaves as an internal negative control for the experiment. Anti-PFA0660w antibodies used in the assay were tested for cross-reactivity with PfHsp70-x-C; no signal was obtained in this blot (Fig. S5A).

**Fig. 3.**
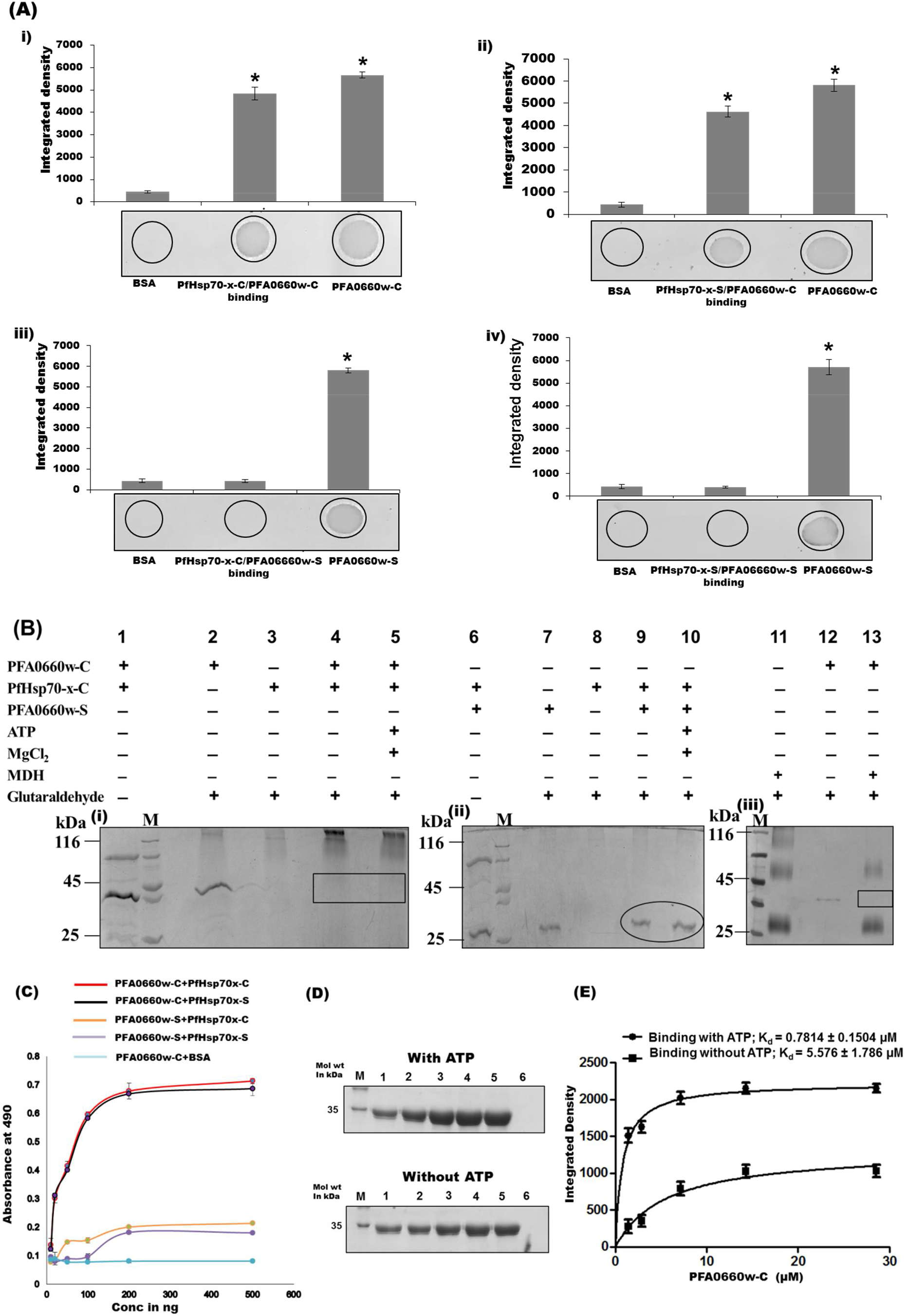
*Interaction of deletion constructs of PFA0660w and PfHsp70-x*. (**A**) *Dot blot assays*. Binding of PFA0660w-C was tested with PfHsp70-x-C (i) and PfHsp70-x-S (ii), and of PFA0660w-S with PfHsp70-x-C (iii) and PfHsp70-x-S (iv). 1μg each of PfHsp70-x constructs were spotted, hybridized with PFA0660w constructs, and probed using anti-PFA0660w antibodies (1:5000). Bar diagram shows plots of intensity measurements from three replicates in an experiment using Image J software. ‘*’ represents statistical significance at p<0.05 relative to BSA. A representative blot from each set is shown. (**B**) *Binding of PFA0660w deletion constructs with PfHsp70-x-C and MDH using glutaraldehyde cross linking*. (i) PFA0660w-C with PfHsp70-x-C (ii) PFA0660w-S with PfHsp70-x-C (iii) PFA0660w-C with MDH. Inset rectangles show desired position of crosslinked PFA0660w-C. Inset circle shows position of cross-linked PFA0660w-S. Lanes M indicates molecular weight marker. (**C**) *Semi-quantitative ELISA*. Concentration-dependent binding curves of PfHsp70-x with PFA0660w where y-axis represents absorbance at 490 nm and x-axis denotes amount of PFA0660w deletion constructs. Error bars represent standard deviation among three replicates. (**D**) *In vitro pull down assays*. Recombinant PfHsp70-x-C was cross-linked to the resin and incubated with increasing concentrations of PFA0660w-C in the presence and absence of ATP to allow binding. Upper panel (with ATP at 1mM) and lower panel (without ATP): Lanes M: Molecular weight marker, lanes 1 to 5: Bound fractions of PFA0660w-C on cross-linked PfHsp70-x-C upon increasing concentrations of PFA0660w-C (1.4286, 2.871, 7.1429, 14.287, 28.5714 μM), lanes 6: Elute from BSA cross-linked beads incubated with PFA0660w-C (negative control). Complete gels are presented in Supplementary Figure 6. (**E**) *Saturation curve for binding of PFA0660w-C with PfHsp70-x-C in the presence and absence of ATP*. Each plotted value represents an average of triplicate determinations with the range indicated by error bars. Calculated K_d_ is indicated on the plot.

The binding of PFA0660w with PfHsp70-x was also studied using glutaraldehyde cross linking assay commonly used to monitor complex formation of Hsp70 with its cochaperones ^41^. Recombinant PFA0660w-C and PFA0660w-S run as monomers on SDS-PAGE when incubated with glutaraldehyde (Fig. 3B lane 2, 7), while PfHsp70-x-C forms oligomers that appear too large to enter the gel (Fig. 3B lane 3, 8). Incubation of PFA0660w-C with PfHsp70-x-C and subsequent crosslinking with glutaraldehyde resulted in the disappearance of the band for PFA0660w-C, suggesting that it binds PfHsp70-x-C (Fig. 3B lane 4, 5). However, band corresponding to PFA0660w-S continued to be observed after crosslinking with PfHsp70-x, indicating their inability to interact (Fig. 3B lane 9, 10). Our results of the glutaraldehyde crosslinking and dot blot assays suggest that binding of PFA0660w with PfHsp70-x is determined by its N-terminal region containing the J domain and G/F region.

Binding of PFA0660w-C was also studied with denatured malate dehydrogenase (MDH – a substrate for chaperones) using glutaraldehyde crosslinking assay (Fig. 3B). MDH alone existed as monomers, dimers and large oligomers upon crosslinking (Fig. 3B lane 11). After PFA0660w-MDH crosslinking, the bands for PFA0660w-C and MDH oligomers disappeared (Fig. 3B lane 13), demonstrating the substrate – chaperone interaction.

PFA0660w-PfHsp70-x binding was also investigated using indirect ELISA assays which reveal significant and specific interaction of PFA0660w-C with both PfHsp70-x constructs in a concentration dependent manner that saturated at higher concentrations (Fig. 3C). We employed *in vitro* pull down assays to measure the binding strength of PFA0660w-PfHsp70-x interaction and to confirm the behaviour of PFA0660w as a co-chaperone of PfHsp70-x. PfHsp70-x-C was cross-linked to amino plus coupling resin and increasing concentrations of PFA0660w-C were added in the presence (substrate interactions abrogated) and absence of ATP (substrate interactions possible). BSA cross-linked beads incubated with PFA0660w-C were used as a negative control. Bound fractions of PFA0660w-C were eluted and separated on SDS PAGE (Fig. 3D), and the integrated density of the bands measured. Binding of PFA0660w-C to PfHsp70-x-C was saturable with an equilibrium dissociation constant (K_d_) of 0.7814 ± 0.1504 μM and 5.576 ± 1.786 μM in the presence and absence of ATP respectively (Fig. 3E). Enhanced binding affinity observed in the presence of ATP suggests the partners to be a chaperone/co-chaperone pair.

*In silico* structural assessment of modelled PFA0660w revealed that its J domain consists of four α helices with the HPD motif residing in the loop region between helices II and III (Fig. 2B). The overall fold of the predicted structure matched that of *E. coli* DnaJ (Hsp40 homolog) (1XBL, 1BQZ, 1BQ0) and human Hdj1 (1HDJ). As reported earlier, the HPD signature motif drives Hsp40-70 binding ^23,42^ while lysine and arginine residues (KR) of the QKRAA motif present in helix IV of the J domain of *E. coli* DnaJ play a crucial role in interaction with DnaK (Hsp70 homolog) ^18^. Multiple sequence alignment of various type II Hsp40 family members recognizes both the HPD motif and ‘KR’ of QKRAA motif to be conserved amongst *Plasmodium* and *non-Plasmodium* species (Fig. S4). Besides, both these are surface exposed on PFA0660w (Fig. 2B), and hence likely to comprise the binding residues in PfHsp40-PfHsp70 interactions.

### Chaperone activity assays

Thermolabile protein substrates have been extensively used to study the chaperone activity of heat shock proteins ^43,44^ Previous studies by Daniyan *et al*. reported additive effect of PFA0660w and PfHsp70-x in suppressing aggregation of the substrate ‘rhodanese’ ^45^. Here, we have employed a set of assays using MDH and beta galactosidase as model substrates for assessing the holdase and foldase functions of recombinant PFA0660w and PfHsp70-x. The holdase function was characterized by quantifying the ability of different combinations of PFA0660w and PfHsp70-x deletion constructs to suppress MDH aggregation (Fig. 4a). The foldase function was investigated by measuring enzymatic activity of beta galactosidase renatured using recombinant Hsps (Fig. 4b).

**Fig. 4.**
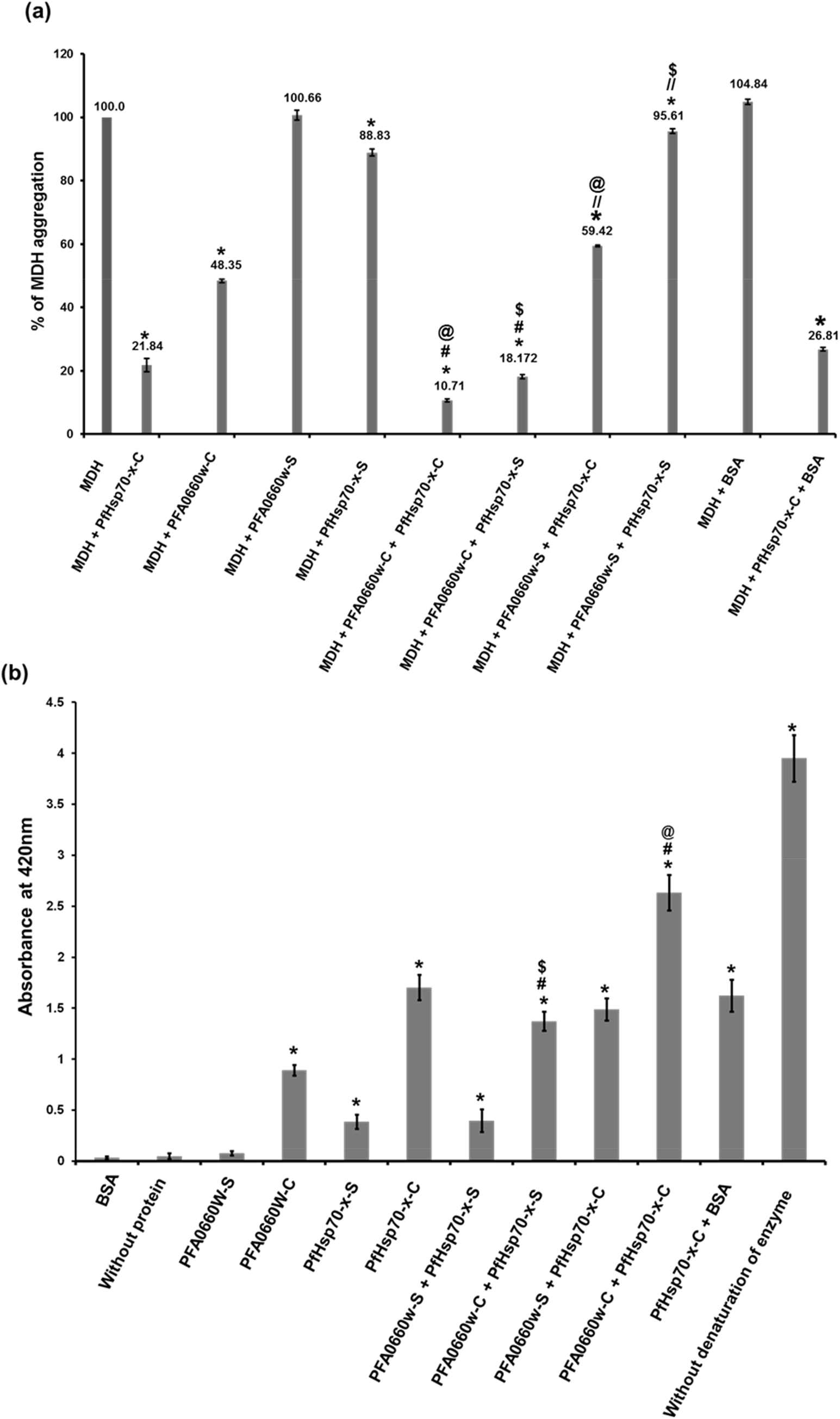
Chaperone activity assays using *deletion constructs of PFA0660w and PfHsp70-x*. (**a**) *MDH aggregation suppression assays*. Percentage of MDH aggregation was observed through absorbance at 360nm in the presence or absence of different combinations of deletion constructs of PFA0660w and PfHsp70-x, as labelled on the plot. MDH aggregation was assumed to be 100% in the absence of chaperone proteins. The average percentage of MDH aggregation is mentioned above each bar; error bars represent the standard deviation. ‘*’, ‘#’, ‘@’, ‘$’, ‘//’ represent statistically significant difference at p<0.05 relative to MDH, PFA0660w-C, PfHsp70-x-C, PfHsp70-x-S and PFA0660w-S respectively. (**b**) *Beta galactosidase refolding assays*. Recovery of activity of denatured beta galactosidase by addition of different deletion constructs of PFA0660w and PfHsp70-x (labelled on the plot) was measured by its ability to convert substrate ONPG to yellow colored ONP whose absorbance was recorded at 420 nm. Experiments were performed in triplicates; error bars represent the standard deviation. ‘*’, ‘#’, ‘@’, ‘$’ represent statistically significant difference at p<0.05 relative to BSA, PFA0660w-C, PfHsp70-x-C and PfHsp70-x-S respectively.

MDH aggregation was monitored spectrophotometrically at 360nm, and considered 100% at saturation (>20minutes) in the absence of any chaperone. PFA0660w-C alone suppressed MDH aggregation efficiently (48.35% aggregation), and was even more effective when added with PfHsp70-x-C (10.71% aggregation) (p<0.05) (Fig. 4a). However, reduction in MDH aggregation using PFA0660w-C and PfHsp70-x-S partnership was relatively less (18.17% aggregation). Interestingly, PFA0660w-S alone or in combination with PfHsp70-x-C or PfHsp70-x-S was unable to suppress MDH aggregation (p>0.05). These data provide evidence that PFA0660w-C having the complete conserved region is able to act as a chaperone independently and can functionally interact with both constructs of PfHsp70-x to reduce MDH aggregation more efficiently, whereas PFA0660w-S is non-functional. Also, PfHsp70-x-C alone reduced MDH aggregation better (21.83% aggregation) than PfHsp70-x-S (88.82% aggregation) (p<0.05) (Fig. 4a), emphasizing the importance of its C-terminal region containing the ‘EEVN’ motif in regulating its chaperone activity. In the absence of MDH, each of the recombinant protein used in the study was found to be thermally stable as none of them aggregated under the assay conditions (Data not shown). BSA was used as a negative control in the experiment to ensure specificity of MDH aggregation suppression by these chaperones.

Beta galactosidase recognizes ONPG as its substrate and cleaves it into ortho-nitrophenol (ONP) and galactose. Absorbance at 420nm measures the amount of ONP produced in the reaction which is a reflection of the ability of chaperones to refold denatured beta galactosidase. In these experiments, the activity of beta galactosidase without denaturation was considered as 100%. BSA itself (negative control) was unable to refold beta galactosidase post denaturation. The activity of denatured beta galactosidase was partially recovered by addition of PFA0660w-C (22.6%), PfHsp70-x-C (43.1%) and PfHsp70-x-S (9.8%) (p < 0.05) but not with PFA0660w-S (p > 0.05) (Fig. 4b). Therefore, PfHsp70-x-C was found to be most efficient in refolding its substrate. Enzymatic activity was significantly increased (66.66%) when PFA0660w-C and PfHsp70-x-C were added together, validating their interaction and suggesting co-ordinated activity (p < 0.05). PFA0660w-C-PfHsp70-x-S partnership was also more effective in refolding action (34.7%, p < 0.05) as compared with PFA0660w-C alone. However, no significant effect was observed when PFA0660w-S-PfHsp70-x-C or PFA0660w-S-PfHsp70-x-S pairs were used in the assay (p > 0.05) (Fig. 4b). BSA-PfHsp70-x-C pair enzyme refolding activity matched that of PfHsp70-x-C alone, verifying the accuracy of the assay (p > 0.05).

Both chaperone activity assays provide evidence for functionality of recombinant constructs of PFA0660w and PfHsp70-x. Inability of PFA0660w-S to suppress MDH aggregation and refold proteins highlights the functional role of PFA0660w’s N-terminal region. Significant difference in activities of PfHsp70-x-C and PfHsp70-x-S suggests the functional importance of its C-terminal region.

### PfHsp70-x interacts with ATS domain of PfEMP1

Since PFA0660w and PfHsp70-x co-localize with PfEMP1 in J-dots ^28,32^, we tested binding of PFA0660w and PfHsp70-x with ATS (acidic terminal segment) domain of PfEMP1 using *in vitro* assays. Preliminary screening was performed using dot blot assays where recombinant PFA0660w-C, PFA0660w-S, PfHsp70-x-C and PfHsp70-x-S were immobilized on nitrocellulose (NC) membrane and allowed to bind with PfEMP1 before probing with ATS specific antisera (Fig. 5A). Both deletion constructs of PfHsp70-x showed significant binding with PfEMP1, whereas no interaction was observed for constructs of PFA0660w (Fig. 5A). Integrated density of each dot further demonstrates significant binding of PfHsp70-x-C and PfHsp70-x-S with ATS domain of PfEMP1 (p<0.05) (Fig. 5A). Absence of signal with immobilized BSA (negative control) demonstrates the specificity of the assay. Anti-ATS antibodies used in the assay were tested for cross-reactivity with recombinant PFA0660w-C and PfHsp70-x-C; no signal was obtained (Fig. S5B). Our semi-quantitative ELISA assays also depict significant binding of PfHsp70-x with ATS domain in a concentration dependent manner in contrast to deletion constructs of PFA0660w (Fig. 5B). Binding of PfHsp70-x-C and PfHsp70-S with PfEMP1 showed saturation at higher concentrations (Fig. 5B). *In vitro* pull down assays followed by western blot analysis also validated the above results. Recombinant PfHsp70-x-C was coupled to aminolink plus coupling resin, and used to pull down recombinant ATS domain of PfEMP1. Elutes from the assay were probed by western blot analysis using anti-ATS antibodies. While a distinct band corresponding to ATS eluted from PfHsp70-x-C coupled beads (Fig. 5C, lane 1; left panel), none was observed for BSA linked beads (negative control) (Fig. 5C, lane 2; left panel). Also, no band was detected when eluted samples were probed with preimmune sera, depicting antibody specificity (Fig. 5C right panel). Bio Layer Interferometry (BLI) was performed to study the binding kinetics of recombinant PfHsp70-x-C with ATS domain of PfEMP1. Fig. 5D (left panel) shows association and disassociation phases of the curves obtained for ATS binding to immobilized PfHsp70-x-C. Equilibrium dissociation constant (K_d_) of 180 ± (1.2× 10^−12^) μM was calculated from the steady state analysis that was obtained by plotting response at equilibrium as a function of analyte concentrations (Fig. 5D, right panel).

**Fig. 5.**
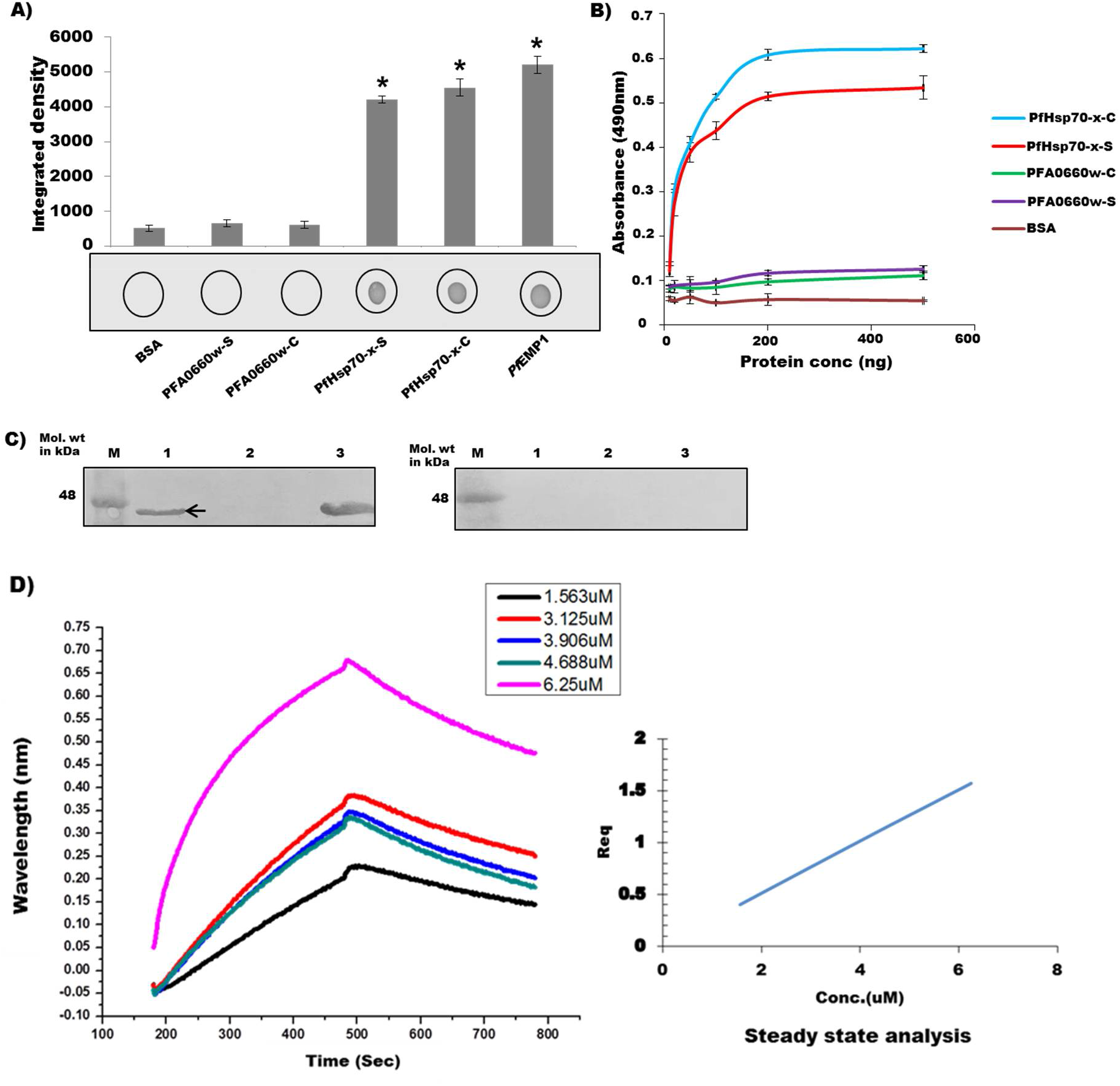
*Interaction of PFA0660w and PfHsp70-x with ATS domain of PfEMP1*. (**A**) *Dot blot assays*. 1μg each of different deletion constructs were spotted, probed with recombinant ATS domain of PfEMP1, and detected using anti-ATS antibodies. Experiments were performed in triplicates. An average of spot intensities was plotted. Error bars show standard deviation. ‘*’ represents statistical significance at p<0.05 relative to BSA (negative control). (**B**) *Semi-quantitative ELISA*. Concentration dependent binding curves were plotted where y-axis represents binding (measured as absorbance at 490nm) and x-axis denotes amount of ATS. Error bars represent standard deviation among three replicates. (**C**) *In vitro pull down assays*. Recombinant PfHsp70-x-C was coupled to amino link coupling resin and incubated with ATS to allow binding. Fraction of ATS bound to PfHsp70-x was eluted and detected by immunoblotting using anti-ATS antibodies (left panel). Right panel represents control blot probed with preimmune sera. Lanes M: molecular weight marker, Lanes 1: Cross linked PfHsp70-x-C incubated with ATS, Lanes 2: Cross linked BSA incubated with ATS, Lanes 3: Recombinant ATS (loading control). Complete blots are presented in Supplementary Figure 6. (**D**) *Biolayer interferometry to study interaction of immobilized PfHsp70-x-C with ATS of PfEMP1*. Association and disassociation curves (left panel) for PfHsp70-x-ATS interaction. Secondary plots (right panel): Steady state analysis shows the net disassociation constant (Kd) of 180 ± (1.2× 10^−12^) μM for PfHsp70-x-ATS interaction.

### C-terminus of PFA0660w binds cholesterol

J-dots are likely membranous structures with unclear lipid composition, and are reported to carry PfHsp40s ^28^. Several chaperones have been reported to localize to specific sub-cellular regions, possibly through binding with lipids ^30,46^. Therefore, we investigated association of chaperone pair ‘PFA0660w-PfHsp70-x’ with common membrane lipids (phosphatidylcholine and cholesterol) using lipid-protein overlay assays ^47^. Binding to a closely related sterol, ‘ergosterol’ and triglycerides were also tested in the assay. Lipids were spotted on NC membrane and incubated with PFA0660w-C, PFA0660w-S, PfHsp70-x-C and PfHsp70-x-S before probing with monoclonal anti-hexahistidine antibodies (Fig. 6A). Our results clearly show the ability of PFA0660w-C and PFA0660w-S to bind specifically with cholesterol but not with other lipids (Fig. 6A left panel). Neither construct of PfHsp70-x showed binding with any of the lipid used in the assay (Fig. 6A left panel). Recombinant protein constructs and BSA were spotted as positive and negative controls respectively. Anti-hexahistidine antibodies used in the assay were tested for cross-reactivity with lipids; no signal was obtained (Fig. S5C). Integrated density of each dot demonstrated significant and specific binding of PFA0660w-C and PFA0660w-S with cholesterol (p<0.05) (Fig. 6A right panel). Additionally, the interaction of PFA0660w with cholesterol could also be seen in semi-quantitative plate-based binding assays where lipids were immobilized and increasing concentration of recombinant proteins were added; PfHsp70-x-C showed no signal (Fig. 6B). Binding saturation was achieved at higher concentrations of PFA0660w-C and PFA0660w-S. Phosphatidylcholine, ergosterol and triglyceride failed to bind either chaperone, defining assay specificity (Fig. 6B). To measure the binding strength of cholesterol with recombinant PFA0660w-C, we used an assay in which the protein was immobilized on Ni-NTA resin before addition of increasing concentrations of cholesterol (Fig. 6C). Bound cholesterol was quantified by Zak’s method ^48^, where binding of cholesterol to PFA0660w-C was saturable with a K_d_ value of 4.82 ± 1.3μM (Fig. 6C). PfHsp70-x-C showed no binding with cholesterol in this experiment. We also performed BLI to study the kinetics of PFA0660w-C-cholesterol interaction. Fig. 6D (left panel) shows association and disassociation phases of the curves obtained for cholesterol binding to immobilized PFA0660w-C. A K_d_ value of 13 ± (7×10^−15^) μM was generated from the steady state analysis which was obtained by plotting response at equilibrium as a function of analyte concentrations (Fig. 6D; right panel).

**Fig. 6.**
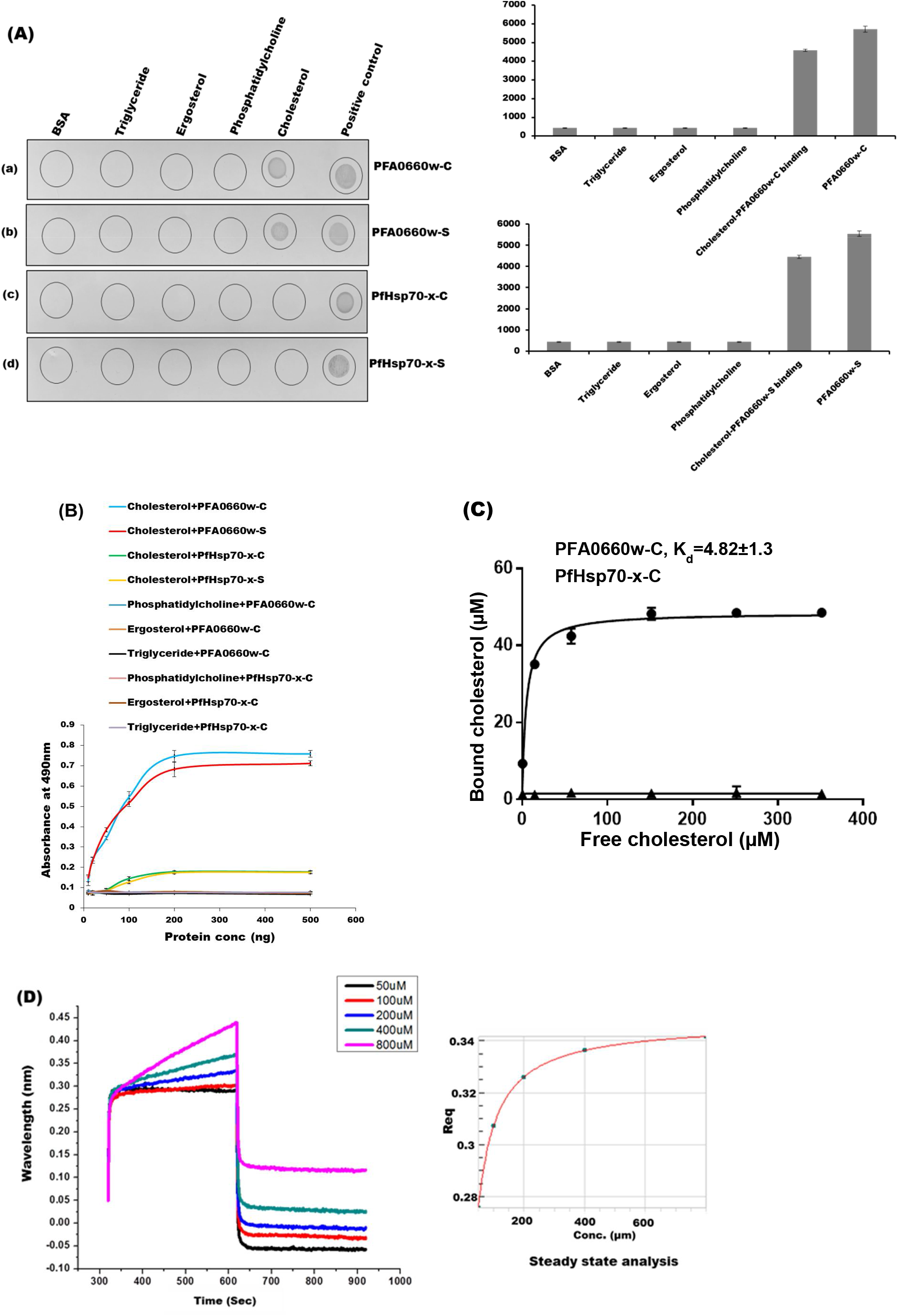
*Interaction of PFA0660w and PfHsp70-x with lipids*. (**a**) *Lipid overlay assays*. Different lipids were immobilized (as labelled above the panel) and deletion constructs were hybridized to the blots (as labelled next to the panel) before probing with monoclonal anti-hexahistidine antibodies. Experiments were performed in triplicates. A representative blot from each set is shown (left panel). Average integrated density from these dots is represented in graphical mode (right panel). Error bars represent standard deviation. ‘*’ represents statistical significance at p<0.05 relative to BSA. (**b**) *Semi-quantitative ELISA*. Concentration dependent binding curves of lipids with different deletion constructs. Lipids were coated and probed with recombinant proteins (as indicated on the plot) followed by monoclonal anti-hexahistidine antibodies. Error bars represent standard deviation among three replicates. (**c**) *Saturation curve for binding of cholesterol with PFA0660w-C using Ni-NTA resin based binding assay*. Each plotted value represents an average of triplicate determinations with the range indicated by error bars. Calculated K_d_ for cholesterol-PFA0660w is indicated on the plot. (**d**) *Biolayer interferometry to study interaction of immobilized PFA0660w-C with cholesterol*. Association and disassociation curves (left panel) for PFA0660w-C-cholesterol interaction. Secondary plots (right panel): Steady state analysis shows the net disassociation constant (K_d_) of 13 ± (7×10^−15^) μM for PFA0660w-cholesterol interaction.

### PFA0660w-S hosts separate binding sites for cholesterol and substrate (MDH)

C-terminal region of PFA0660w carries both substrate and cholesterol binding sites. We analyzed the surface representation of modelled C-terminal PFA0660w to understand where cholesterol is likely to bind on this protein, and found two major hydrophobic grooves (Fig. 7A). One constituted the substrate binding pocket ^38^, while the second was present on the extreme C-terminal end of this domain (Fig. 7A). The latter is reported to form the dimer interface in Hsp40 homologs from other species ^38,39^. Therefore, we investigated whether denatured MDH and cholesterol bind to separate sites or compete for binding to PFA0660w-S. PFA0660w-S bound Ni-NTA beads were saturated with denatured MDH before incubation with increasing concentrations of cholesterol to test whether this causes displacement of bound MDH. SDS-PAGE analysis of bound fractions shows no effect of cholesterol addition on MDH band intensity, suggesting existence of distinct cholesterol and substrate binding pockets on PFA0660w-S (Fig. 7B, upper panel). Absence of leaching of PFA0660w-S bound MDH in unbound fractions upon cholesterol addition validates these findings (Fig.7A, lower panel). Direct binding of denatured MDH non-specifically to Ni-NTA beads was experimentally ruled out (Fig. 7B, lane 7, upper panel). Densitometric analyses of band intensities of PFA0660w-S and MDH observed on SDS-PAGE was performed to quantify the result (Fig. 7B, right panel).

**Fig. 7.**
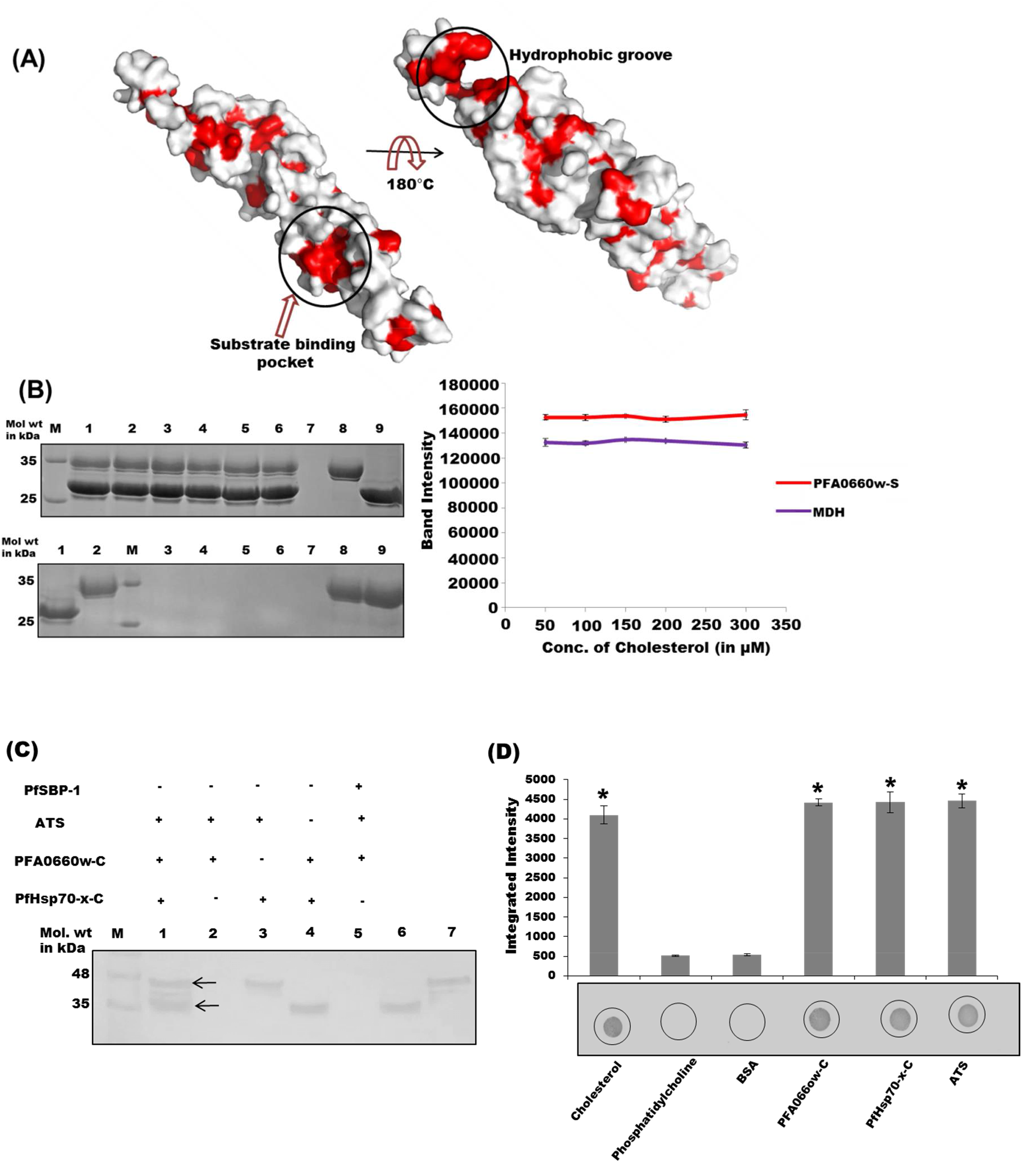
*Cholesterol/MDH binding assays and complex binding assays*. (**A**) *Surface representation of C-terminal region of PFA0660w*. Substrate binding pocket and major hydrophobic groove are encircled. Hydrophobic residues are marked in red. (**B**) *Cholesterol and MDH binding on PFA0660w-S bound Ni-NTA beads*. Upper panel: Lane M: Molecular weight marker, lanes 1 to 5: (bound) fractions of PFA0660w-S – MDH complex with increasing cholesterol concentrations (50, 100, 150, 200, 300 μM), lane 6: bound PFA0660w-S –MDH with no cholesterol (positive control), lane 7: elute from MDH addition to beads in the absence of bound PFA0660w-S (negative control), lane 8: purified MDH, lane 9: purified recombinant PFA0660w-S. Lower panel: Unbound fractions (leached MDH) on addition of cholesterol. Lane 1: purified MDH, lane 2: purified recombinant PFA0660w-S, Lane M: Molecular weight marker, lanes 3-7: Flow throughs (unbound fractions) from incubated cholesterol in increasing concentrations (50, 100, 150, 200, 300 μM respectively), lane 8: unbound MDH from positive control, lane 9: unbound MDH from negative control. Graph showing measured band intensity of PFA0660w-S and MDH on addition of increasing cholesterol concentration (right panel). Experiment was performed in triplicates, and an average of intensities plotted. Complete gels are presented in Supplementary Figure 6. (**C**) *PFA0660w-PfHsp70-x-ATS complex formation*. Lane M: Molecular weight marker, lane 1: Cross linked PfHsp70-x-C incubated with PFA0660w and ATS, lane 2: cross linked PFA0660w-C incubated with ATS (negative control), lane 3: Cross linked PfHsp70-x-C incubated with ATS (positive control), lane 4: cross linked PfHsp70-x-C incubated with PFA0660w-C (positive control), lane 5: cross linked PfSBP-1 incubated with PFA0660w-C and ATS (negative control), lane 6: purified recombinant PFA0660w-C, lane 7: purified recombinant ATS. Complete blot is presented in Supplementary Figure 6. (**D**) *Complex formation of Cholesterol, PFA0660w, PfHsp70-x and ATS*. Lipids/ proteins (labelled) were immobilized, and allowed to bind PFA0660w-C followed by PfHsp70-x-C and then ATS before probing with anti-ATS antibodies. Experiment was performed in triplicates, and an average of integrated density of each dot was plotted as a graph (upper right panel). ‘*’ represent statistical significance at p<0.05 relative to negative controls. A representative blot is shown (bottom right panel).

### PfHsp70-x binds PFA0660w and PfEMP1 simultaneously *in-vitro* to form a complex

Since PfHsp70-x binds both PFA0660w and ATS of PfEMP1, we tested its potential to attach with both proteins simultaneously to form a complex using *in-vitro* aminoplus coupling resin based pull down assays ^36^. Recombinant PfHsp70-x-C was coupled to the resin, and allowed to bind PFA0660w-C, followed by ATS. Elutes from the assay were analysed by western blot analysis using specific polyclonal antisera against PFA0660w and ATS simultaneously. Distinct bands of ATS and PFA0660w-C eluted from PfHsp70-x-C cross linked beads were observed (Fig 7C, lane 1), while no band was detected where beads were crosslinked with PFA0660w-C and incubated with ATS (lane 2). Cross linked PfHsp70-x-C incubated with ATS and PFA0660w-C separately showed PfHsp70-x-C– ATS and PfHsp70-x-C–PFA0660w-C interactions (positive controls; lane 3 and lane 4). PfSBP-1 (Pf Skeleton binding protein experimentally shown to not bind ATS) was cross linked to the coupling resin as a negative control in this experiment (lane 5) ^26^.

To test association of this complex with lipids, we performed lipid-protein overlay assays. Cholesterol and phosphatidylcholine were immobilized on NC membrane and allowed to bind PFA0660w-C, PfHsp70-x-C and ATS sequentially. The blot was probed with specific polyclonal antisera against ATS. A strong signal was detected for Cholesterol-PFA0660w-C-PfHsp70-x-C-ATS, but none for immobilized phosphatidylcholine (Fig. 7D). PfHsp70-x-C–ATS and PfHsp70x-C–PFA0660w-C (positive controls) and BSA (negative control) were included in the assay.

## Discussion

Parasite encoded exported chaperones are involved in the cellular refurbishment of infected human erythrocytes, and play an important role in the establishment of protein folding pathways in the host cell. Understanding the molecular interplay of Pf heat shock proteins provide functional insights into malaria biology. In this study, we have explored various aspects of PFA0660w-PfHsp70-x interaction, and attempted to delineate its function in infected red blood cells.

Two deletion constructs each of PFA0660w and PfHsp70-x were cloned in pET-28a(+) vector and overexpressed in soluble form in a bacterial expression system [BL21 (DE3) *E. coli* cells]. A recent study reported expression of recombinant hexahistidine-tagged PFA0660w in the insoluble fraction when cloned in pQE30 vector ^45^. Biochemical characterization and atomic resolution three dimensional structures of Hsp40s from other organisms (yeast Sis1, human Hdj1) reveal these to exist as dimers ^38,39^. Deletion of Sis1 dimerization unit affects its ability to function as a chaperone ^38^. Here, the cleft formed between the two monomers helps in holding the substrate peptide and deliver it to Hsp70 ^38,39^. C-terminal regions of both *C. parvum* Hsp40 (cgd2_1800) ^40^ and *E. coli* DnaJ play a role in their dimerization, and the latter is reported to be necessary for its chaperone function ^49^. In contrast to these studies, we found PFA0660w to exist as a monomer when investigated by gel permeation chromatography and glutaraldehyde crosslinking assays (Fig. 2A, 3B). Our results corroborate those of Daniyan *et al*., who had previously reported refolded PFA0660w to attain monomeric conformation in solution ^45^. Lack of PFA0660w dimer formation is also consistent with a recent report on the characterization of a novel Type III Hsp40, ‘Tbj1’ from *Trypanosoma brucei* ^50^. Our *in silico* analysis using multiple sequence alignment and dimer interface residue mapping on homology models predicts the role of hydrophobic interactions in homo-dimerization of Hsp40s. Since oligomeric state of Hsp40 proteins is directly linked to their function, we emphasize that PFA0660w is structurally and functionally different from its counterparts in other organisms, including humans. Such striking structural differences between host and parasite protein counterparts identify essential ‘PFA0660w’ as a candidate for structure-based drug design against Pf caused malaria.

Our binding studies reveal that the PFA0660w-PfHsp70-x interaction is mediated by the N-terminus of PFA0660w containing its J domain and the G/F region (Fig. 3), which may serve to provide specificity to this molecular interaction. Independent ability of PfHsp70-x-S to bind with PFA0660w-C recognises the importance of the ATPase domain on PfHsp70-x in forming the major binding site.

Daniyan *et al*. have previously reported functionality of recombinantly expressed PFA0660w and PfHsp70-x, and an additive effect in chaperone activity when used together ^45^. Interrogating the role of different domains in chaperone action is vital for understanding the functional partnership of the PFA0660w - PfHsp70-x pair. Our chaperone activity assays using various deletion constructs reveal that conserved length construct PFA0660w-C could function alone, and showed cooperative action with both constructs of PfHsp70-x (Fig. 4). However, PFA0660w-S (C-terminal region) was inactive in either assay, highlighting the functional significance of its N-terminus. These results are coherent with previous literature where Hsp40 proteins have been shown to exhibit intrinsic chaperone activity besides acting as co-chaperones ^44,51^. Similarly, significant difference in the capability of PfHsp70-x-S and PfHsp70-x-C to perform their chaperone activity identifies the importance of its C-terminal region carrying the ‘EEVN’ motif in regulating its chaperone function. ‘EEVD’, a regulatory motif found in eukaryotic Hsp70s at the extreme carboxyl terminal has been reported to effect its ATPase activity and binding to both its partner ‘Hsp40’ and substrates ^52^. Collectively, our domain analysis identifies N-terminus of PFA0660w and C-terminus of PfHsp70-x as critical for functionality of this chaperone pair.

PfHsp70-x deletion from parasites leads to delayed export of PfEMP1 to iRBC surface ^34^. Both PFA0660w and PfHsp70-x have been observed to localize to specialized mobile structures in the erythrocyte cytosol, termed ‘J-dots’ ^28,32^. These are different from the well characterized Maurer’s clefts and are believed to be formulated with a distinct lipid composition ^28^. Also, cholesterol rich microdomains have been implicated in the trafficking of major cytoadherence ligand ‘PfEMP1’ ^29^. In light of the above facts, we tested binding of PFA0660w and PfHsp70-x with ATS domain of PfEMP1, and identified *in vitro* interaction of PfHsp70-x with ATS (Fig. 5). PfHsp70-x-S showed similar binding as the conserved length protein, suggesting its N-terminal region to contain the binding site for ATS. This is the first report of direct binding of PfHsp70-x with ATS domain of PfEMP1, which adds another piece in solving the jigsaw puzzle of PfEMP1 trafficking.

Kulzer *et al*. had indicated tight association of PFA0660w-GFP with J-dots, possibly by protein-cholesterol interaction ^28^. However, *in vivo* trials to understand PFA0660w-cholesterol interaction in J-dots did not provide conclusive evidence, probably owing to cholesterol concentrations below detectable limits of the method used. Therefore, we tested direct binding of PFA0660w and PfHsp70-x with cholesterol and other common membrane lipids. Our data demonstrate specific interaction of PFA0660w with cholesterol, while PfHsp70-x did not show any lipid binding properties (Fig. 6). PFA0660w-S having only its C-terminal region also showed similar cholesterol binding pattern, suggestive of a role for its carboxyl terminus in anchorage to J-dots. Recently, a report on targeting of GFP tagged PFE0055c (another J-dot resident Hsp40 protein) to infected erythrocytes showed that its SBD fused with a signal peptide and PEXEL motif on the N-terminus is necessary and sufficient for recruiting the protein to J-dots ^53^. Since C-terminal region of PFA0660w shares 54% identity with PFE0055c, we hypothesize the role of its SBD in targeting PFA0660w to J-dots by a direct cholesterol dependent interaction. We further propose that PfHsp70-x may be recruited to J-dots through its specific interaction with PFA0660w, though involvement of other molecular players is not ruled out.

We marked the hydrophobic residues on modelled C-terminal PFA0660w (Residues 226-398) to identify a probable binding site for cholesterol on this protein. *In silico* analysis of its surface representation revealed the presence of an additional far C-terminal major hydrophobic groove apart from its substrate binding pocket. This region is ordinarily involved in homo-dimer formation in Hsp40 counterparts from other species ^38,39^. Considering the monomeric state of PFA0660w, it is likely that this additional pocket may form its cholesterol binding site (Fig. 7A). Our *in vitro* Ni-NTA based assays revealed presence of distinct binding sites for denatured MDH (substrate) and cholesterol on C-terminus of PFA0660w (Fig. 7B). This highlights the ability of PFA0660w to dually function as a chaperone and membrane linking molecule simultaneously.

Since PfHsp70-x binds both PFA0660w and ATS of PfEMP1, we tested whether both recombinant proteins bind PfHsp70-x together to form a complex, or compete with each other for attachment to PfHsp70-x. Our assays clearly show that PfHsp70-x binds both PFA0660w and PfEMP1 simultaneously *in vitro* suggestive of separate binding sites for these on PfHsp70-x (Fig. 7C). Proteome analysis coupled with biochemical studies revealed PfHsp70-x to exist in different high molecular weight complexes at J-dots, and the parasitophorous vacuole (PV) ^33^. Though co-immunoprecipitation experiments failed to detect PfEMP1 in these complexes probably due to experimental limitations, presence of PfEMP1 here is still very likely ^33^. Our *in vitro* studies also suggest that PfEMP1 may associate with PfHsp70-x and Hsp40 containing complexes. Further, our dot blot assay shows that PFA0660w-PfHsp70-x-PfEMP1 complex is able to link itself to cholesterol via PFA0660w (Fig. 7D). Our results are in line with previous studies by Frankland *et al*. who demonstrated the role of cholesterol rich structures in export of PfEMP1 to the surface of infected erythrocytes ^29^. It is interesting to speculate here that cholesterol depletion may interrupt linkage of PFA0660w-PfHsp70-x-PfEMP1 complex to J-dots, and prevent subsequent delivery of PfEMP1 to iRBC membrane. There also exists evidence for formation of large chaperone containing soluble complexes of PfEMP1 to mediate its trafficking across the host cytosol ^11,54^. From our studies and previous data, we hypothesise a model for the role of PFA0660w-PfHsp70-x chaperone pair in PfEMP1 transport via direct association with cholesterol containing J-dots (Fig. 8). Our model is supported by the observation of all three members of the complex in J-dots ^28,32^. We propose formation of this complex either pre or post exit from PV to erythrocyte cytosol through the translocon. Since PfHsp70-x has been dually localized to both PV and J-dots ^32^, it is probable that PEXEL negative PfHsp70-x delivers cargo (e.g. PfEMP1) to the erythrocyte cytoplasm with the assistance of PEXEL positive proteins like PFA0660w. A recent report has detailed the association of PfHsp70-x with PTEX on the trans side of the PVM ^33^. After crossing the parasite confines, the said complex may insert into the J-dots before being delivered to the iRBC surface either via Maurer’s clefts or directly to the knobs.

**Fig. 8.**
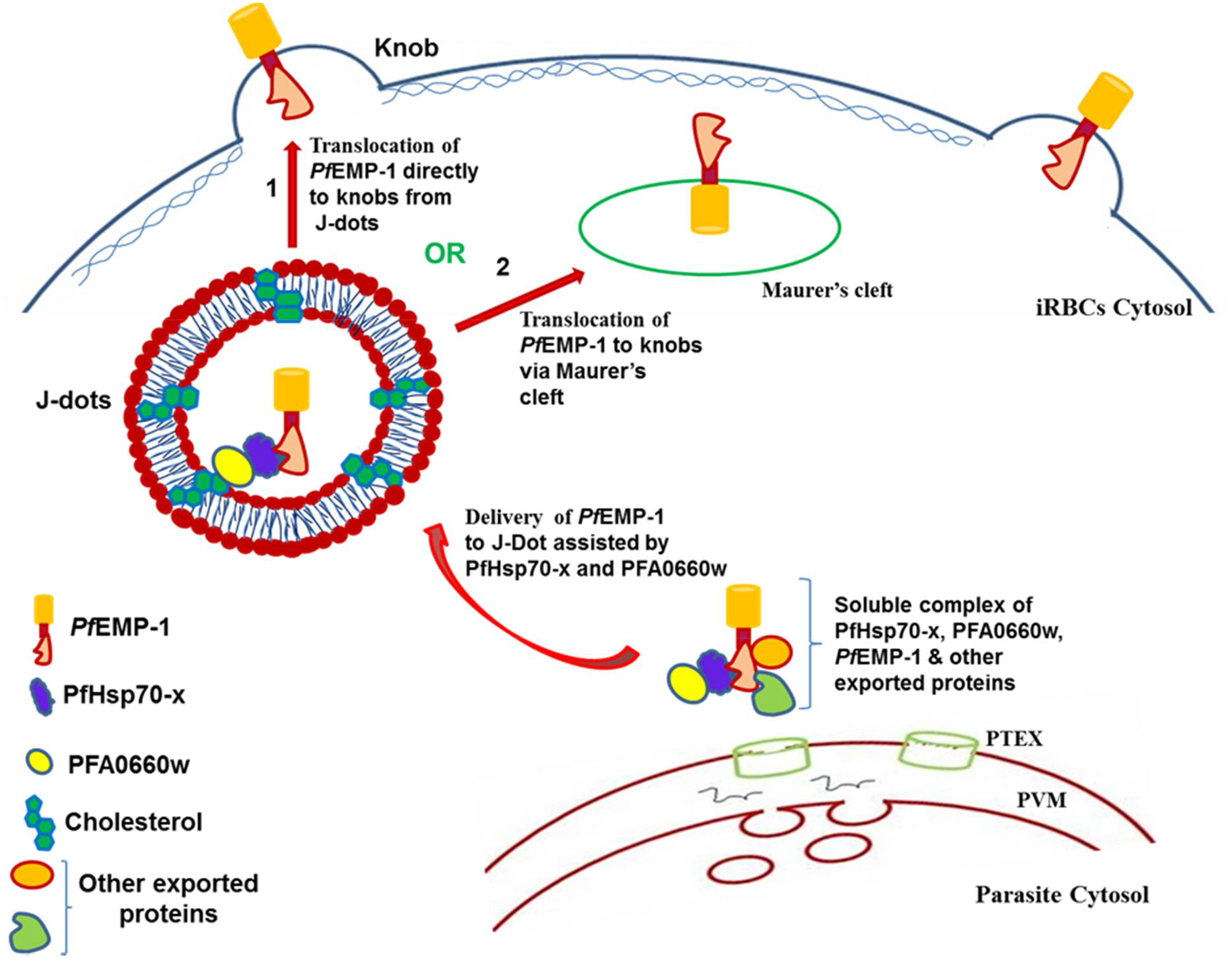
*A hypothetical model depicting the probable function of PFA0660w-PfHsp70-x chaperone pair*. Soluble complexes of PfEMP1 with PFA0660w-PfHsp70-x chaperone pair may be delivered to J-dots via binding of PFA066w with cholesterol. PFA0660w-PfHsp70-x-PfEMP1 complex could be either contained inside of the J-dots or be linked on their exterior. J-dots may then serve to deliver PfEMP1 to the iRBC surface either via Maurer clefts or by direct translocation to the knobs.

Emerging resistance to the currently available drugs against malaria is a major threat to mankind, and demands development of newer agents to combat this disease ^55,56^. Understanding molecular interactions of essential Pf proteins may provide the key to identification of new inhibitors through structure-based drug design. Since PFA0660w is essential for parasite survival and is significantly different from its host counterpart, drugs that target its molecular networking with PfHsp70-x or cholesterol are likely to impact malaria pathogenesis and parasite survival.

## Material and methods

### Cloning, expression and purification of recombinant proteins

DNA corresponding to deletion constructs of PFA0660w and PfHsp70-x were PCR amplified, and cloned in pET-28a(+) bacterial expression vector before expression in *E. coli* BL21 (DE3) cells. Recombinant proteins were purified using Ni-NTA affinity chromatography followed by gel permeation chromatography. Gel filtration chromatography was performed using AKTA prime plus on Superdex 200 10/300 GL column (GE Healthcare). Purified PFA0660w-S was used for raising polyclonal antibodies in rabbits commercially (Merck, Bangalore, India).

To confirm protein identity, purified recombinant protein constructs of PFA0660w and PfHsp70-x were resolved on SDS-PAGE and transferred on nitrocellulose (NC) membrane. NC membrane was blocked overnight at 4°C in 5 % BSA. Following washing, the blot was probed with HRP linked monoclonal anti-hexahistidine antibody (1:2000) for 2 hours, and developed with DAB/H_2_O_2_ substrate.

Recombinant ATS domain of PF08_0141 was purified for experiments as previously described ^36,37^.

### Structure prediction and validation

Blastp against the RCSB Protein Databank (PDB) was used to find suitable templates for modelling the J domain and C-terminal region of PFA0660w ^57^. Models were generated using Swiss Model tool ^58^ and were refined by 3D refine, a protein structure refinement server ^59^. The refined models were evaluated using VERIFY 3D, ERRAT and RAMPAGE programs ^60–62^. PyMol was used for visualization and analysis of protein structures ^63^. Protein Interactions Calculator (PIC) server was used to identify the possible dimer interface residues of *C. parvum* Hsp40 (PDB ID: 2Q2G) and yeast Sis1 ^64^.

### Dot blot assays

For screening protein-protein interactions of PfHsp70-x with PFA0660w, and various Hsp constructs with ATS of PfEMP1, the recombinant purified PfHsp70-x or Hsp proteins (~1μg each) were immobilized on NC membrane. Blots were blocked with 5% BSA in PBS for 2 hrs, and hybridized with ~20μg of probe protein (PFA0660w or ATS of PfEMP1 respectively) for 1 hr. After extensive washing, blots were incubated with probe protein specific antibodies (1:5000) for 1 hr followed by horseradish peroxidise (HRP)-conjugated goat anti-rabbit (for PFA0660w) or HRP conjugated goat anti-mice (for ATS) secondary antibody (1:2000). Relevant recombinant purified Hsp proteins were used as positive controls, while BSA was used as a negative control in the assay. Integrated densities of dots were measured using ImageJ after background correction ^65^.

### Glutaraldehyde cross linking

Recombinant proteins (~60μg) were suspended in crosslinking buffer (50mM sodium phosphate pH 8, 100mM NaCl in the presence or absence of ATP and MgCl_2_ (2mM each)) in a total volume of 100μl, and incubated at 37°C for 15 minutes. 5μl of 2.3% glutaraldehyde was added to the reaction and incubated at 37°C for another 30 minutes. MDH was denatured at 48°C prior to crosslinking with PFA0660w-C. The reaction was terminated by adding 10μl of stop solution (1 M Tris-HCl, pH 8.0), and samples were resolved on 12% SDS-PAGE.

### Plate based interaction studies

*In vitro* interactions of PFA0660w with PfHsp70-x, and various Hsp constructs with ATS of PfEMP1 were examined by indirect ELISA. 100ng each of purified bait proteins (recombinant purified PfHsp70-x or Hsp proteins) were coated on ELISA plates and blocked with 5% BSA in PBS overnight at 4°C. The coated ligands were incubated with increasing concentrations of prey proteins (PFA0660w or ATS of PfEMP1 respectively; range: 10ng to 500 ng) for 2 hrs at room temperature, followed by extensive washing with 1X PBS. Washed plates were incubated with anti-PFA0660w-S (1:5000) or anti-ATS antibodies (1:5000) followed by anti-rabbit or anti-mice HRP conjugated secondary antibodies (1:10,000) for two hrs. Plates were developed using 1mg/ml OPD (o-phenylenediamine dihydrochloride) containing H2O2, and absorbance measured at 490nm.

For investigating protein-lipid interaction, a similar protocol as described above was followed where 100ng of lipids were coated on ELISA plates and allowed to bind with varying concentration of hexahistidine tagged recombinant proteins (PFA0660w-C, PFA0660w-S, PfHsp70-x-C and PfHsp70-x-S). Bound proteins were detected by incubating plates with monoclonal anti-hexahistidine-HRP conjugated antibodies (1:10,000).

### Protein pull down assay

Pierce chemical co-immunoprecipitation kit was used according to the manufacturer’s protocol to evaluate direct protein-protein interaction of PfHsp70-x (bait protein) with either PFA0660w or ATS of PfEMP1 (prey). Briefly, 10μl of coupling resin was cross-linked with 20μg PfHsp70-x-C for 2 hrs. After extensive washing, protein coupled resin was incubated with increasing concentrations of PFA0660w-C (1.4286, 2.871, 7.1429, 14.287, 28.5714 μM) in the presence and absence of 1mM ATP or 50 μg of ATS respectively diluted in 200 μl of binding buffer. Unbound protein was removed by repeated washing with binding buffer, followed by elution of prey protein. As a negative control, BSA was coupled to the resin as bait before incubation with PFA0660w-C or ATS. Samples of bound PFA0660w-C were resolved on 12% SDS-PAGE and band intensities measured using ImageJ after background correction. *Kd* was determined by fitting the curve with nonlinear regression using GraphPad Prism 7.0 software (GraphPad Software, CA, USA). Eluted samples incubated with ATS were subjected to western blot analysis using anti-ATS antibodies.

### Malate dehydrogenase (MDH) aggregation suppression assays

MDH aggeregation suppression assays were carried out as previously reported ^44^.

### Beta galactosidase refolding assays

50μl of a 1mg/ml solution of beta galactosidase (sigma) was denatured by incubation with equal volume of denaturation buffer (25mM HEPES pH 7.5, 50mM KCL, 5mM MgCl_2_, 5mM beta-mercaptoethanol and 6M guanidine-HCl) at 30°C for 30 minutes. Denatured substrate enzyme was refolded by treatment with 1ml refolding buffer (25mM HEPES pH 7.5, 50mM KCl, 5mM MgCl_2_, 10mM DTT, 1mM ATP) containing 1μM of recombinant protein constructs at 37°C for 30 minutes. Enzyme activity was tested by addition of 200μl substrate ‘o-nitrophenyl-β-D-galactoside’ (ONPG: 4 mg/mL in 0.1M phosphate buffer pH 7) for 10 minutes followed by termination with 300μl of 1M Na_2_CO_3_ as stop solution. BSA was used in place of recombinant proteins as a negative control. Absorbance was recorded at 420nm for each reaction.

### Biolayer interferometry assays

Protein–protein and protein-lipid interaction studies were carried out using Forte Bio Octet K2 instrument at 25°C in solid black 96-well plates. The binding assay kinetics were carried out with PFA0660w-C and PfHsp70-x-C as ligands. 300 μg/ml of different His tagged ligands (PFA0660w-C or PfHsp70-x-C) were used to load the surface of Ni-NTA biosensors for 200s, and the sensor washed for 300s. Different analyte concentrations of ATS domain of PfEMP1 (1.563, 3.125, 3.906, 4.688, 6.25 μM) or cholesterol respectively (50, 100, 200, 400, 800 μM) were used for the binding studies in a total volume of 200 μl/well. Systematic baseline drift correction was carried out using assay buffer. Data analysis and curve fitting were determined using Octet Data analysis software version 9.0, and curve fitting analysis performed with the 1:1 interaction binding model. Global analysis of the complete data was carried out using nonlinear least squares fitting to find a single set of binding parameters at all the tested analyte concentrations, and steady-state kinetic analyses carried out for all data.

### Lipid protein overlay assays

Nitrocellulose membranes were soaked in TBS buffer before 0.5μg of methanol suspended lipids were spotted and allowed to dry for 1 hr at room temperature. Membrane was blocked with 5 % BSA in TBS for 2 hrs followed by incubation with 25μg of hexahistidine tagged proteins (PFA0660w-C, PFA0660w-S, PfHsp70-x-C and PfHsp70-x-S). Membranes were washed with 0.05% TBST followed by TBS and incubated with monoclonal anti-hexahistidine-HRP conjugated antibodies (1:2000) for 2 hrs, and developed.

This assay was also employed to test complex formation ability of cholesterol-PFA0660w-C-PfHsp70-x-C-ATS. A similar protocol as described above was followed except that after immobilization of lipids, the membrane was hybridized with PFA0660w-C followed by PfHsp70-x-C and then ATS (1 hr each). Blots were incubated with anti-ATS antibodies (1:5000) for 1 hr followed by HRP-conjugated goat anti-mice secondary antibody (1:2000), and developed.

### Cholesterol Binding assay

100μg of recombinant protein was allowed to bind with Ni-NTA resin followed by blocking with 5% BSA in TBS. After washing, increasing concentration of cholesterol (10-400 μM in binding buffer (50mM Tris, pH 7.4, 150mM NaCl, 0.004% NP-40)) was allowed to bind with protein immobilized resin for 4 hrs. Resin was washed with binding buffer, and cholesterol bound protein was eluted with elution buffer (binding buffer + 250mM imidazole). Concentration of bound cholesterol was estimated by Zak’s method ^48^ using a standard curve for cholesterol. Concentration of free cholesterol was calculated by subtracting bound concentration from total cholesterol added in the assay. *Kd* was determined by fitting a hyperbola directly to the saturation isotherm using GraphPad Prism 7.0 software (GraphPad Software, CA, USA).

### Bead based cholesterol/MDH binding assay

Ni-NTA resin bound to PFA0660w-S was used to test whether MDH and cholesterol bind this Hsp40 together to form a complex, or compete for binding. 20μg of purified recombinant PFA0660w-S was allowed to bind with Ni-NTA beads followed by blocking with 5% BSA in TBS. Bound PFA0660w-S was saturated with a standardized fixed concentration of denatured MDH (200μg) for 2 hrs with intermittent mixing; MDH was incubated at 48°C for denaturation prior to binding. After extensive washing with TBS, increasing concentrations of cholesterol (50, 100, 150, 200, 300μM) were added and incubated for 4 hrs on a rocker. Flow-through post cholesterol binding would show displaced MDH (unbound sample), if any. Resin bound proteins (PFA0660w-S - MDH) were eluted with elution buffer containing imidazole and resolved on 12% SDS-PAGE. Band intensities were measured using ImageJ ^65^.

### Complex binding assay

*In-vitro* formation of a complex between PFA0660w, PfHsp70-x and ATS was tested using Pierce co-immunoprecipitation kit. 25μg of PfHsp70-x-C was coupled to the beads, and blocked with 5% BSA overnight at 4^°^C. Coupled resin was incubated with recombinant PFA0660w-C (2 hrs) followed by ATS for 2 hrs. PfHsp70-x-C was cross linked and incubated with ATS and PFA0660w-C separately as positive controls. PFA0660w-C and PfSBP-1 were coupled to beads separately and incubated with ATS and PFA0660w-C respectively as negative controls. Eluted samples were subjected to western blot analysis to probe for PFA066w-C and ATS by anti-PFA066w-C and anti-ATS antibodies respectively on the same blot.

### Statistical analysis

Statistical significance was analysed by one way ANOVA (Tuckey’s post hoc test) using SPSS version 16 ^66^. Values with *P* value < 0.05 were considered as statistically significant.

## Acknowledgements

Laboratories of PCM and RH were funded by Department of Biotechnology (DBT), Govt. of India. AB was a DBT-Senior Research Fellow (Department of Biotechnology-Govt. of India). VK was initially funded by DST, and is now a UGC National Fellow (UGC, Govt. of India).

## Author Contributions

AB^1^: Experimental design, experimentation, data analysis and manuscript writing. VK: Conducted complex binding assay and some pull down assays, manuscript preparation. AB^3^: Conducted the BLI experiments. JJP: Planned and executed the BLI experiments and analysed BLI data. RH: Experimental design, data analysis and manuscript writing. PCM: Conception of idea, experimental design, data analysis and manuscript writing.

## Abbreviations

iRBC: Infected red blood cell
Pf: *Plasmodium falciparum*
Hsp40: Heat shock protein 40
Hsp70: Heat shock protein 70
PfEMPl: *Plasmodium falciparum* erythrocyte membrane protein1
ATS: Acidic terminal segment
BSA: Bovine serum albumin
MDH: Malate dehydrogenase
IPTG: Isopropyl-1-thio-β-D galactopyranoside
ONPG: o-nitrophenyl-β-D-galactoside
ONP: ortho-nitrophenol
Ni-NTA: Nickel-nitrilotriacetic acid beads
PBS: Phosphate-buffered saline
HRP: Horseradish peroxidase
SDS–PAGE: Sodium dodecyl sulphate– polyacrylamide gel electrophoresis
NBD: Nucleotide-binding domain
SBD: Substrate binding domain

